# Conserved and newly acquired roles of PIF1 homologs in tomato (*Solanum lycopersicum*)

**DOI:** 10.1101/2021.10.29.466498

**Authors:** Miguel Simon-Moya, M Victoria Barja, Luca Morelli, Daniele Rosado, Linlin Qi, Gianfranco Diretto, Tomás Matus, Briardo Llorente, Jaime F. Martinez-Garcia, Alain Goossens, Magdalena Rossi, Manuel Rodriguez-Concepcion

## Abstract

PHYTOCHROME INTERACTING FACTORS (PIFs) are transcription factors that interact with the photoreceptors phytochromes and integrate multiple signaling pathways related to light, temperature, defense and hormone responses. PIFs have been extensively studied in *Arabidopsis thaliana*, but less is known about their roles in other species. Here, we investigate the role of the two homologs of PIF1 found in tomato (*Solanum lycopersicum*), namely PIF1a and PIF1b. Analysis of gene expression showed very different patterns, indicating a potential evolutionary divergence in their roles. At the protein level, light regulated the stability of PIF1a, but not PIF1b, further supporting a functional divergence. Phenotypic analyses of CRISPR-Cas9-generated tomato mutants defective in PIF1a or PIF1b or both revealed conserved and newly acquired roles compared to Arabidopsis PIF1. Both PIF1a or PIF1b were found to regulate seed germination, photosynthetic pigment biosynthesis and fruit production. However, only PIF1a-defective mutants showed defects on root hair elongation, flowering time and fruit growth and softening. We did not identify any process altered only in plants lacking PIF1b. Together, these data show that neofunctionalization has taken place in tomato, illustrating the potential of these transcription factors to acquiring new roles in different species.

## INTRODUCTION

Light is the main source of energy for plants but also a key source of environmental information that strongly influences plant developmental processes such as seed germination, phototropism, seedling development, chloroplast biogenesis and floral induction, among others. Plants are equipped with photoreceptors that detect and transduce light signals to trigger such responses. Phytochromes (phys) are probably one of the best known photoreceptors in plants, with five members (phyA to phyE) found in the model plant *Arabidopsis thaliana*. They are specialized in detecting and transducing light in the red (R) and far-red (FR) wavelengths. Phytochromes exist in two photoconvertible forms, an inactive R light-absorbing Pr form and an active FR light-absorbing Pfr form. The inactive Pr form is located in the citosol. Following the R light-mediated conversion, the Pfr form translocates to the nucleus where it can interact with a family of transcription factors called Phytochrome-Interacting Factors (PIFs). This interaction causes PIF phosphorylation and inactivation, eventually triggering a rapid and global transcriptional reprogramming (Park et al., 2012; Qiu et al., 2017; Park et al., 2018).

PIFs are members of the basic helix-loop-helix (bHLH) transcription factor superfamily (Ni et al., 1998). The Arabidopsis PIF family is composed of 8 members. All members have an Active phyB Binding domain (APB) but only PIF1 and PIF3 present an additional Active phyA Binding domain (APA) in their sequence (Leivar and Quail, 2011). Arabidopsis PIFs have been proposed as central integrators of light and other signaling pathways related to environmental (e.g. temperature and defense) and endogenous (e.g., hormone) cues (REF). Besides their role as transcription factors directly regulating the expression of thousands of genes, PIFs also interact with different types of regulatory proteins to modulate plant responses. (Leivar and Quail, 2011). In particular, Arabidopsis PIF1 is a repressor of seed germination, seedling photomorphogenesis (including chloroplast development and photosynthetic pigment - chlorophylls and carotenoids-accumulation) and flowering (Toledo-Ortiz et al., 2010, 2014; Bou-Torrent et al., 2015; Chenge-Espinosa et al., 2018; Huq et al., 2004; Oh et al., 2004, 2006, 2007; Wu et al., 2018). PIF1 also functions in leaves, repressing carotenoid gene expression to adjust the levels of these important photoprotectors in response to environmental (i.e. light and temperature) signals (Toledo-Ortiz et al. 2010; Toledo-Ortiz et al. 2014). Similar to other PIFs, however, the role of PIF1 in other plant species remains much less studied.

In tomato (*Solanum lycopersicum*), two homologs of PIF1 have been reported, namely PIF1a (Solyc09g063010) and PIF1b (Solyc06g008030) (Llorente et al., 2016b, Rosado et al., 2016). PIF1a was proposed to repress carotenoid biosynthesis in green fruit, when the presence of chlorophylls prevents most R light to enter the fruit and hence causes a self-shade effect that promotes the accumulation of PIFs (Llorente et al., 2016a, 2016b). During ripening, chlorophylls degrade, resulting in lower PIF levels and derepressed production of carotenoid pigments that provide the red color of ripe fruit. Strikingly, *PIF1a* expression did not decrease but increased during ripening, a mechanism that was interpreted as a ‘gas-and-brake’ effect providing a more robust control of fruit carotenoid biosynthesis (Llorente et al., 2016a, 2016b).

Consistent with the negative role on the regulation of carotenoid biosynthesis reported for the members of the so called PIF quartet (PIFQ: PIF1, PIF3, PIF4 and PIF5) in Arabidopsis (Toledo-Ortiz et al., 2010, 2014; Bou-Torrent et al., 2015), silencing of the genes encoding PIF1a or PIF4 in tomato resulted in higher carotenoid levels (Llorente et al., 2016; Rosado et al., 2019). PIF4 was also found to be a regulator of thermomorphogenesis and inductor of leaf senescence and flowering in both Arabidopsis (Sakuraba et al., 2014; Song et al., 2014; Zhang et al., 2015) and tomato (Rosado et al., 2019). On the other hand, tomato PIF3 was identified as a regulator of steroidal glycoalkaloid (SGA) biosynthesis, a cholesterol-derived family of compounds that are not present in Arabidopsis but and contribute to pathogen defense in Solanaceae (Wang et al., 2018). Together, these results suggest that both evolutionary conservation and neofunctionalization occurs within the PIF family when comparing Arabidopsis and tomato. Here we aimed to further explore this possibility by focusing on PIF1 homologs.

## RESULTS

### Tomato PIF1 homologs show different expression profiles

As an initial step to investigate whether the two PIF1 homologs present in tomato might play different physiological roles, we analyzed the expression pattern of both *PIF1a* and *PIF1b* genes using the Bio-Analytic Resource for Plant Biology (BAR, University of Toronto). According to these data (Fig. 1A), both *PIF1a* and *PIF1b* are expressed at similar levels in two organs: roots (where they both are low expressed) and leaves (where their transcript levels are much higher). Levels of *PIF1a* transcripts are higher than those of *PIF1b* in all stages of flower and fruit development. *PIF1b* expression increases as fruit grows until it reaches its final size at the mature green (MG) stage. During ripening, *PIF1b* expression does not change but *PIF1a* expression exhibits a strong increase as fruits move from the breaker (BR) to the red ripe (RR) stage. This result is in accordance with previously reported experimental data (Llorente et al., 2016b; Rosado et al., 2016). We next analyzed the gene co-expression networks (GCN) of *PIF1a* and *PIF1b* as an additional way of testing whether these two transcription factors might be involved in similar processes. We used data from TomExpress (Abdullah et al., 2017) to get a list of the 500 most highly co-expressed genes with *PIF1a* and *PIF1b* in three different organs: root, leaf and fruit (Supp Fig 1). Strikingly, we found virtually no overlapping of the GCNs of *PIF1a* and *PIF1b* in roots and leaves, whereas only about 10% of genes showed either positive (40 genes) or negative (62 genes) co-expression with both *PIF1a* and *PIF1b* in fruit (Supp Fig 1).

**Fig 1.**
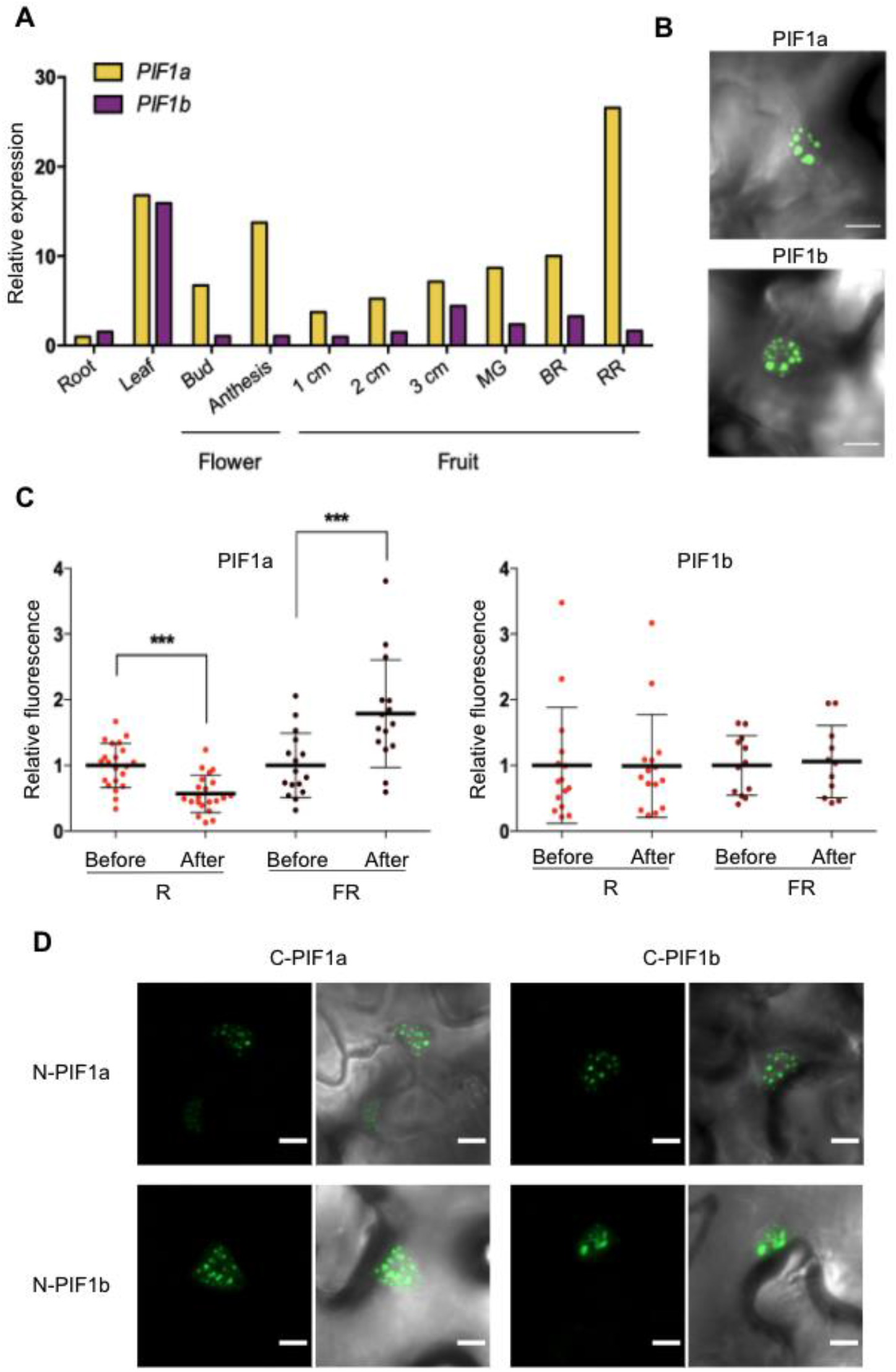
Expression profiles and protein features of PIF1a and PIF1b. A. Levels of PIF1 a and PIF1 b transcripts in different plant organs. Data are obtained from BAR (University of Toronto) and expressed relative to the lowest expression level found in the study (i.e. PIF1a expression in roots). B. Confocal microscopy images of GFP fluorescence in the nucleus of N. benthamiana leaf cell transiently expressing PIF1a or PIF1 b fused to GFP. Scale bar = 10 μm C. Quantification of GFP fluorescence of nuclei from leaf areas. Error bars indicate SD of at least 15 different nuclei. Asterisks mark statistically significant changes in student’s test (*** = p < 0.001) D. BiFC analysis of PIF1a and PIF1b protein-protein interaction.Confocal microscopy images of GFP fluorescence in N. benthamiana leaf cells transiently expressing the indicated proteins fused to N- or C-terminal GFP halves for BiFC analysis. Images of representative nuclei showing either GFP fluorescence alone (left) or overlapped with bright field images (right) are shown for every combination. Scale bar =10 μm.

### A mutation in the APB domain of PIF1b impacts ligh-dependent stability but does not prevent heterodimerization with PIF1a

Beyond expression patterns, protein stability is a key factor in the regulation of PIF activity. PIF1a and PIF1b are very similar in their amino acid sequence (primary structure), showing characteristic basic Helix-Loop-Helix (bHLH) and active phyB-binding (APB) domains like all PIFs plus an active phyA-binding (APA) domain that in Arabidopsis is only present in PIF1 and PIF3. However, the APB-binding domain of PIF1b presents an amino acid substitution that changes a conserved Q residue to G (Rosado et al., 2016). It was proposed that this sequence change could lead to a disrupted interaction with phyB and hence a differential light-dependent degradation compared to PIF1a (Bu et al., 2011b; Park et al., 2012). In order to experimentally confirm whether PIF1a and PIF1b abundance is differentially controlled by light, we used C-terminal GFP-tagged versions of these proteins transiently expressed in *Nicotiana benthamiana* leaves. Both PIF1a-GFP and PIF1b-GFP proteins were localized in nuclear bodies (Fig. 1B) (Gramegna et al., 2018), as expected for PIF transcription factors (Al-sady et al., 2006; Shen et al., 2005; Trupkin et al., 2014). Once we confirmed that the GFP-tagged proteins were correctly localized, we irradiated the agroinfiltrated leaf with either R or FR light for 30 min and took pictures of the same area before and after irradiation (Fig. 1C and Supp Fig 2). PIF1a-GFP fluorescence signal decreased after the exposure to R and increased after FR, a dynamic behavior that is typical of a PIF which interacts with phyB after a prolonged illumination (Shen et al., 2008). In contrast, PIF1b-GFP abundance was not altered by R or FR exposure (Fig. 1C and Supp Fig 2). It is therefore likely that, as predicted, the single amino acid change present in the APB domain of PIF1b prevents normal interaction with active phyB and hence disrupts its light regulation, likely impacting the biological activity of the protein.

Although it is not predicted that the mutation in the APB domain could interfere with the interaction of PIF1b with other proteins besides phyB, we decided to test it by taking advantage of the reported capacity of PIFs to interact with each other forming homodimers and heterodimers (Toledo-ortiz et al., 2003; Bu et al., 2011a). Using Bimolecular Fluorescence Complementation (BiFC), we confirmed that PIF1a and PIF1b are able to interact in the nucleus, more specifically in nuclear bodies, forming homodimers but also PIF1a-PIF1b heterodimers (Fig. 1D and Supp Fig 3).

### Single and double mutants were obtained using a CRISPR/Cas9 strategy

In order to unveil the processes in which PIF1 homologs were involved, we generated knock-out lines in the tomato MicroTom variety using the CRISPR/Cas9 technology. We selected lines with a single nucleotide insertion for *PIF1a* and a two-nucleotide deletion for *PIF1b*. Both mutations caused a frameshift predicted to result in shorter PIF1a and PIF1b proteins that lacked the APA and bHLH domains, likely resulting in non-functional mutants (Supp Fig 4). We also generated double mutants and carried out multiple serial crosses with untransformed wild type (WT) plants to remove the CRISPR/Cas9 transgene and possible off-targets. We next used the resulting lines (*pif1a, pif1b* and *pif1a pif1b*) to test whether the loss of function of PIF1a, PIF1b, or both, interfered with the expression of other genes encoding PIFQ homologs (Supp Fig 5). The qPCR results showed no significant differences in transcript levels between WT and mutant leaves and MG fruit, with the only exception of decreased *PIF1a* transcript levels in PIF1a-defective mutants (Supp Fig 5).

### Both PIF1a and PIF1b repress germination under FR light

Arabidopsis PIF1 is an essential player in light-regulated seed germination. Under FR light, PIF1 accumulates in the nucleus and represses the germination process, while in the presence of R light, the activated phys degrade PIF1 and germination takes place (Oh et al., 2004; Seo et al., 2009; Shinomura et al., 1994; Lee et al., 2012; Shi et al., 2013). Similar to Arabidopsis, germination of tomato seeds is repressed by FR light and promoted by R light (Shichijo et al., 2001; Appenroth et al., 2006; Auge et al., 2009). To test whether tomato PIF1 homologs might be involved in the light-dependent regulation of seed germination, we exposed seeds from WT and mutant lines to hourly pulses of R or FR light as previously described (Appenroth et al., 2006). Treatment with R light resulted in similar germination rates for WT and mutant *pif1a*, *pif1b* and *pif1a pif1b* seeds (Fig. 2A). By contrast, FR light treatment repressed germination of WT seeds, as expected, but had little impact on the germination of mutant seeds (Fig. 2A). This indicates that both PIF1a and PIF1b repress seed germination in tomato under FR light, similar to that observed for PIF1 in Arabidopsis.

**Fig 2.**
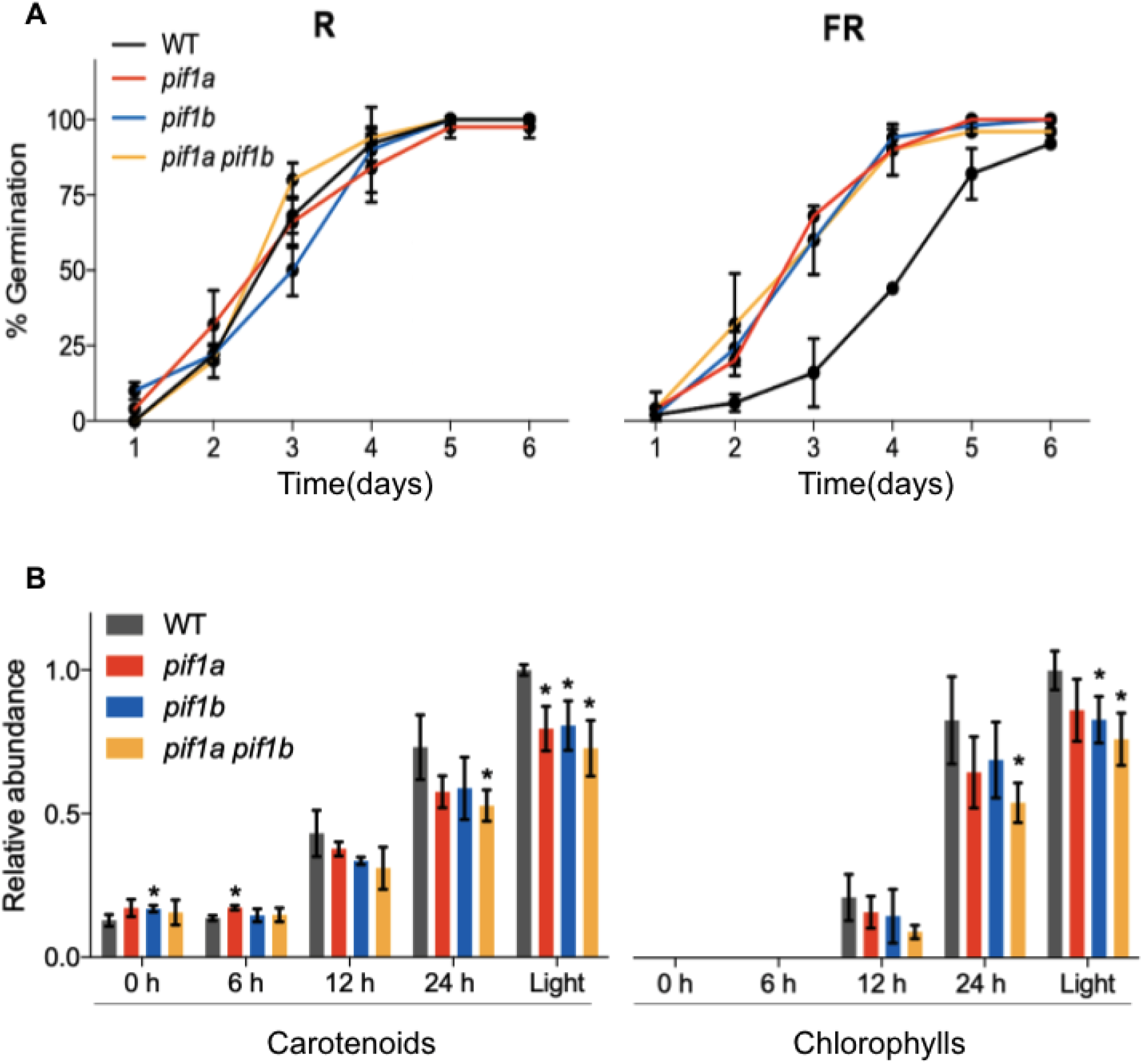
Germination and de-etiolation assays. A. Kinetics of germination of the 4 indicated genotypes under hourly pulses of R (left) or FR (right) for 6 days. Error bars indicate SD of 2 biological replicates with at least 50 seeds each. B. Carotenoid and chlorophyll levels during de-etiolation. WT and mutant seedlings were germinated in the dark for 7 days and then illuminated with white light for the indicated times. Seedlings germinated and grown under continuous white light for 7 days were used as controls. Carotenoid and chlorophyll levels are represented relative to those in light-grown WT seedlings. Error bars indicate SD of at least 3 biological replicates. Asterisks mark statistically significant changes in the indicated genotype compared to the WT according to t-student test (* = p < 0.05).

### PIF1a and PIF1b do not repress but induce photosynthetic pigment production

Arabidopsis PIF1 was shown to negatively regulate carotenoid and chlorophyll biosynthesis during seedling de-etiolation (Huq et al., 2004; Moon et al., 2008; Toledo-Ortiz et al., 2010). PIF1 accumulates in dark-grown (i.e., etiolated) seedlings, directly repressing the expression of genes supporting the production of photosynthetic pigments and the differentiation of chloroplasts. In the dark-to-light transition, PIF1 is degraded, allowing rapid biosynthesis and accumulation of chlorophylls and carotenoids and the transformation of etioplasts into chloroplasts (Toledo-Ortiz et al., 2010, 2014; Bou-Torrent et al., 2015). To test whether tomato PIF1 homologues contributed to carotenoid and chlorophyll production during seedling de-etiolation, we germinated WT and mutant seeds in the dark for 7 days, and then illuminated them with white light for an additional day. Samples were collected at different times after illumination for measuring photosynthetic pigment levels by HPLC. Plants germinated and grown in the light were used as controls. In stark contrast with the results reported in Arabidopsis, the absence of PIF1a, PIF1b, or both, in tomato seedlings led to a deceleration in the light-triggered accumulation of carotenoids and chlorophylls (Fig. 2B). A similar phenotype has been reported in PIF-defective Arabidopsis mutants when they were left for too long in the dark (Monte et al., 2004). In the case of tomato, however, reduced levels of photosynthetic pigments were also detected in mutant seedlings that were germinated and growth for 7 days in the light (Fig. 2B), suggesting that the tomato PIF1 homologs might not be repressors but activators of chlorophyll and carotenoid biosynthesis in photosynthetic tissues.

### Root hair elongation is blocked only in PIF1a-defective lines

While doing the germination and de-etiolation experiments, we noticed that tomato single *pif1a* and double *pif1a pif1b* mutants had hairless roots (Fig. 3A), a phenotype that has never been reported in PIF-deficient Arabidopsis mutants. We confirmed that Arabidopsis *pifq* mutants defective in PIF1 but also in the other members of the PIF quartet showed no obvious defects in root hair development (Fig. 3B). Scanning Electron Microscope (SEM) analysis of tomato roots showed that root hair primordia were indeed present in *pif1a* and *pif1a pif1b* roots, suggesting that the loss of PIF1a activity does not interfere with root hair initiation but prevents elongation (Fig. 3C). This phenotype was very robust, affecting all the seedlings in homozygous populations of single *pif1a* and double *pif1a pif1b* mutants, and being absent from WT and *pif1b* mutants. Therefore, the results suggest that PIF1a, but not PIF1b, is required for root hair elongation in tomato.

**Fig 3.**
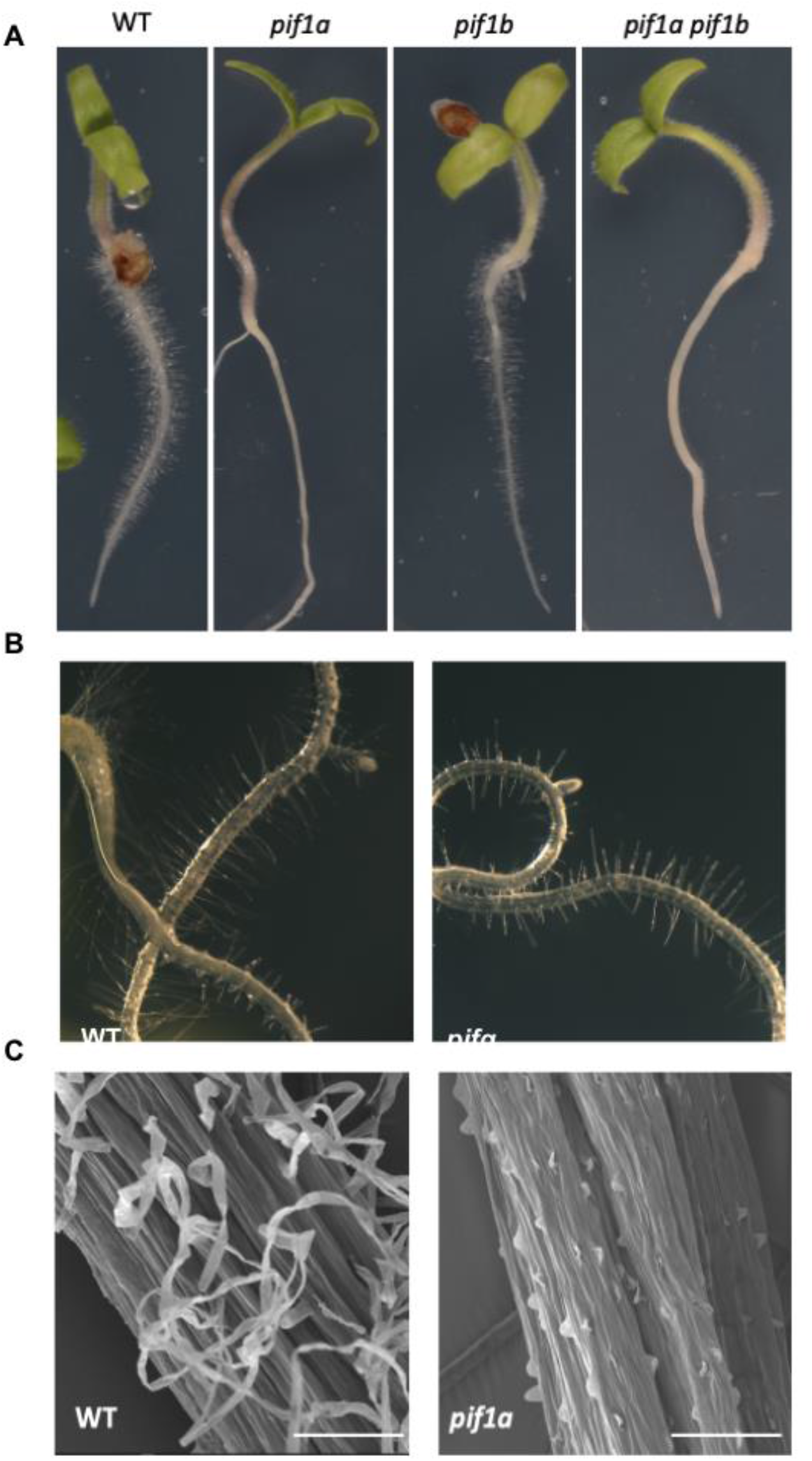
Root hair development in *pif* mutants. A. Visual phenotype of WT and *pif1* mutant seedlings. B. Arabidopsis roots of WT and *pifq* mutant. C. SEM images of tomato WT and *pif1* mutant roots. Scale bar = 100 μm

### Loss of PIF1a might result in early flowering

Arabidopsis *pif1* mutants show early flowering and up-regulated expression of the major flowering-promoting genes (Wu et al., 2018). We assessed the flowering time of tomato *pif1a* and *pif1b* mutants by measuring two different parameters: (1) the number of leaves produced when the first flower reached anthesis and (2) the number of days from sowing to anthesis. The first parameter (number of leaves) indicated a slight but statistically significant reduction of flowering time (i.e. early flowering) in single *pif1a* and double *pif1a pif1b* mutants, but not in *pif1b* plants (Supp Fig 6). By contrast, the second parameter (number of days) showed no differences between genotypes (Supp Fig 6). A number of other studies in the literature report differences in flowering when using one of the evaluated parameters but not when using the other (Calvert, 1959; Giliberto et al., 2005).

### Fruits defective in PIF1 homologs show normal carotenoid and ethylene production

Despite the obvious differences between tomato and Arabidopsis fruits, many of the genes known to be involved in the control of tomato fruit development and ripening are homologous to Arabidopsis genes controlling different developmental processes (Itkin et al., 2009; Gapper et al., 2013; Seymour et al., 2013; Pesaresi et al., 2014). In particular, genes identified to participate in light signaling in Arabidopsis were later found to be involved in fruit development and ripening in tomato (Monte et al., 2004; Bianchetti et al., 2018; Lupi et al., 2019). Using our tomato mutants defective in PIF1a and PIF1b, we aimed to investigate possible roles of these two homologs in different processes associated with tomato fruit development.

PIF1a was previously found to repress **carotenoid biosynthesis during ripening** (Llorente et al., 2016b). Fruits from transgenic lines with a partially silenced *PIF1a* gene were found to accumulate higher levels of carotenoids when ripe (Llorente et al., 2016b), and a similar phenotype has been recently reported in *PIF4*-silenced fruits (Rosado et al., 2019). The levels of carotenoids but also other related isoprenoid metabolites regulated by PIFs (chlorophylls and tocopherols) were quantified by HPLC in WT and mutant fruits at three different ripening stages, from MG to RR. Strikingly, virtually no differences were found in the levels of any of these metabolites between WT and mutant fruit in any of the stages analyzed (Supp Fig 7). Consistently, fruit pigmentation was normal in all mutant lines. Carotenoid production and many other processes associated with fruit ripening are regulated by the hormone ethylene (Giovannoni, 2004). Because an important crosstalk between PIFs and many plant hormones (including ethylene) has been described in Arabidopsis (Lau and Deng, 2010; de Lucas and Prat, 2014), we next tested whether ethylene metabolism was affected during fruit ripening of the tomato PIF1-defective mutant lines by GC-MS (Pereira et al., 2017). We found no significant differences between genotypes (Supp Fig 7), concluding that none of the tomato PIF1 homologs regulates ethylene production during fruit ripening.

### Tomato fruit size, weight and texture are regulated by PIF1a but not PIF1b

Down-regulation of PIF4 leads to a decrease in fruit yield and size (Rosado et al., 2019). In order to test whether the tomato PIF1 homologs might also participate in the regulation of fruit yield and size, we quantified the number and volume of fruits produced by mutant lines. Interestingly, *pif1a* mutant plants produced less fruits than WT controls, while no differences were found in the case of *pif1b* or *pif1a pif1b* lines (Fig. 4A). Fruit size was estimated by measuring the volume of groups of 10 fruits harvested from the plant at the RR stage. Opposite to that observed in the case of PIF4-silenced fruits (Rosado et al., 2019), *pif1a* and *pif1a pif1b* fruits were found to be bigger than WT or *pif1b* fruits (Fig. 4B). We next measured the weight of 100 individual RR fruits from each genotype and confirmed that single *pif1a* and double *pif1a pif1b* mutant plants developed not only bigger but also heavier fruits (Fig. 4C). Furthermore, we confirmed that the difference in weight in PIF1a-defective fruits was maintained after drying the fruits (Supp Fig 8), which means that it is not due to increased water content but results from an enhanced accumulation of dry matter.

**Fig 4.**
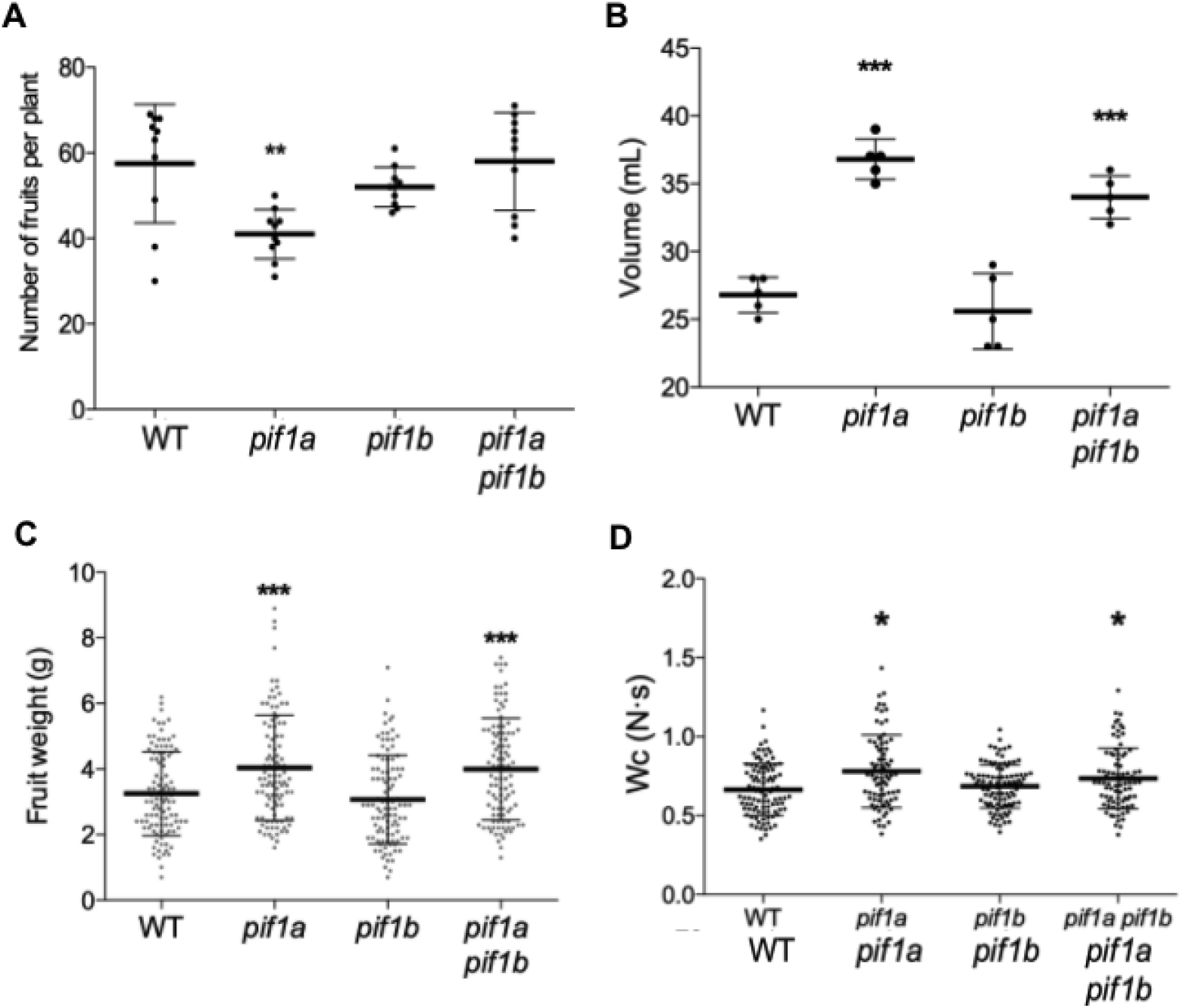
Fruit traits in *pif1* mutants. A. Fruit yield in tomato pif1 mutants. Plot represents the total number of fruits produced by individual 19-week-old plants grown on soil in the greenhouse under long day conditions. B. Fruit volume in tomato pif1 mutants. Plot represents the volume of 5 groups of 10 fruits each. Fruits were collected at the RR stage from plants grown on soil in the greenhouse under long day conditions. C. Weight of 100 individual RR fruits of each genotype. Error bars indicate SD. D. Fruit texture in tomato pif1 mutants. Plot represents Wc measurements (compression tests) of 100 individual RR fruits of each genotype. Error bars indicate SD. Asterisks mark statistically significant changes in the indicated genotype compared with WT according to one-way ANOVA (* = p < 0.05, ** = p < 0.01, *** = p < 0.001)

The observation that *pif1a* mutants produced more and bigger fruits prompted us to investigate whether the texture of ripe PIF1a-deficient fruits was also altered. In order to measure fruit hardness, we used a texture analyzer fitted with 50 mm plate probe to perform a compression test (Kabas and Ozmerzi, 2008; Camps and Gilli, 2017). This test provides a texture parameter called Wc, which is the mechanical work needed to reach a 5% deformation of the fruit. Higher Wc values were obtained in the case of *pif1a* and *pif1a pif1b* mutants (Fig 4D), indicating a higher resistance to deformation (i.e., harder fruit).

## DISCUSSION

### PIF1a and PIF1b work together to regulate seed germination, photosynthetic pigment biosynthesis and fruit production

Several lines of evidence support the conclusion that duplicated copies of the *PIF1* gene in tomato have functionally diverged. The first one is the differential protein stability of PIF1a and PIF1b under different light conditions. Our data suggest that the amino acid substitution in the APB-binding domain of PIF1b likely leads to a failure in the interaction with phyB (Fig. 1). This result suggests that during evolution, PIF1b may have lost its capacity to transduce phyB-mediated light signals. Contrasting with this, PIF1b-defective lines showed light-dependent phenotypes during seed germination (Fig. 2A) and seedling de-etiolation (Fig. 2B). It has been reported that phyB regulates the R light-induction and FR light-inhibition of seed germination when using single light pulses (Appenroth et al., 2006). In contrast, phyA would be involved in the induction and inhibition when the pulses were applied hourly (as we did in our experiment). Therefore, the lack of interaction with phyB would not be a problem to transduce the phyA-dependent light signal mediating the germination response under a multi-pulse experimental system. The involvement of phyA and other phys besides phyB in the control of photosynthetic pigment accumulation during seedling deetiolation has been also described (Su et al., 2015). In fact, phyA has been described as the photoreceptor with a major contribution to initiating photomorphogenesis in Arabidopsis (Casal et al., 2003; Seo et al., 2004). This predominant role of phyA would also explain why PIF1b might regulate this process despite being unable to interact with phyB.

Light-dependent seed germination and photosynthetic pigment biosynthesis during seedling de-etiolation are regulated by both PIF1a and PIF1b proteins (Fig. 5). In fact, loss of any of the two PIF1 homologs produces the same phenotype than the lack of both in double mutants (Fig. 2A and 2B). This made us think that both PIF1a and PIF1b are part of the same signal transduction pathway. We speculate that there might be a threshold amount of PIF1 activity needed to trigger the signaling pathways required to inhibit germination or to accumulate photosynthetic pigments during seedling development. In WT lines, both PIF1a and PIF1b are required to reach this threshold. If one of the genes is lost, the remaining amount of PIF1 activity would not be enough to normally trigger the process. Loss of all tomato PIF1 activity in double *pif1a pif1b* mutants would have the same effect observed in single mutants, i.e. when the threshold is not met by losing only one of the two genes. Another possible scenario based on the BiFC result (Fig. 1D) is that maybe PIF1a and PIF1b need to form a heterodimer to regulate the expression of the genes involved in seed germination and photosynthetic pigment accumulation during de-etiolation. So, when one of the elements of this complex is missing, these processes are affected (Fig. 5). This complex would be interacting with phyA through both of the elements or with phyB through just PIF1a.

**Fig 5.**
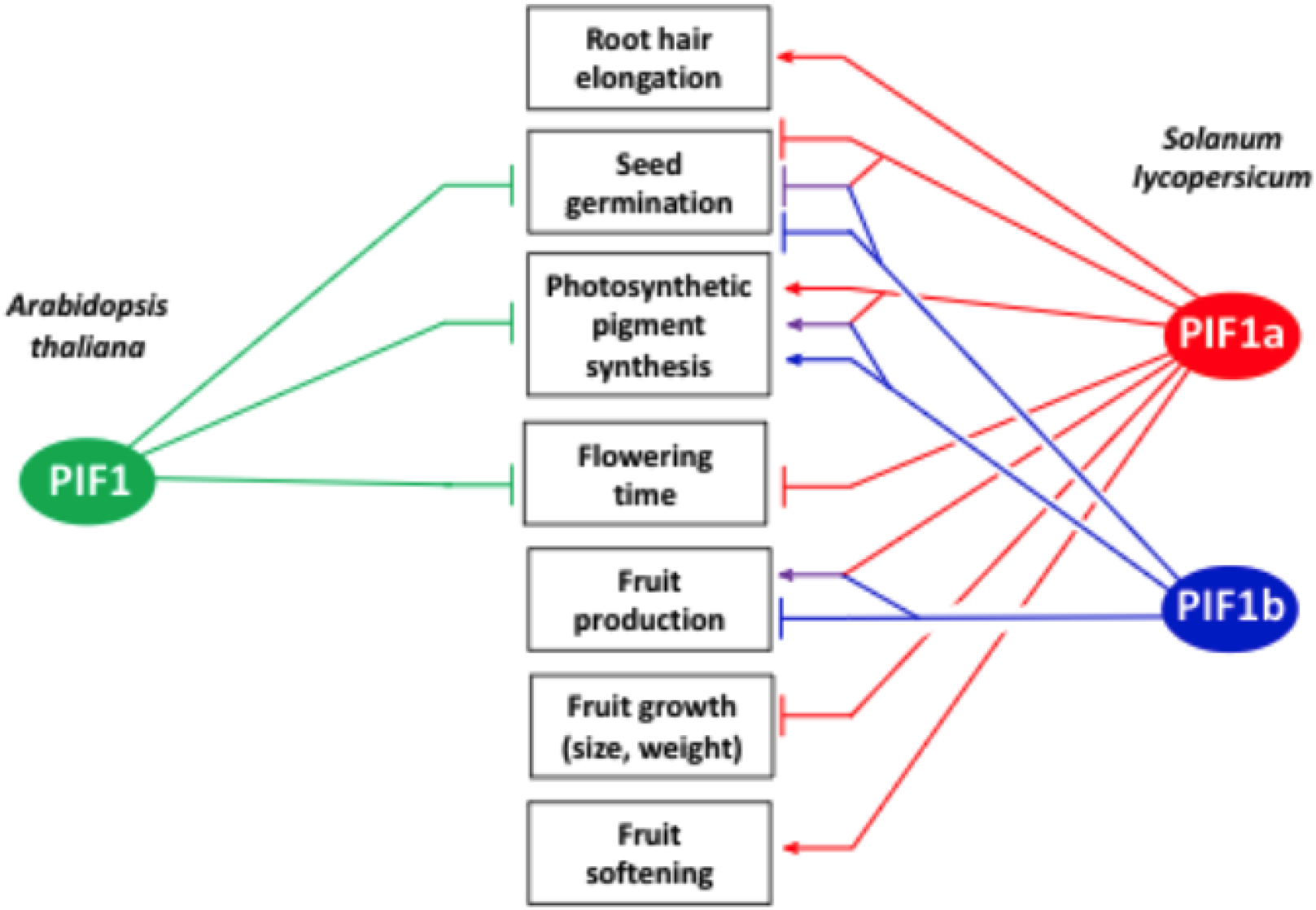
Schematic summary of proposed roles for PIF1 homologs in Arabidopsis and tomato. Arrows represent induction and bars represent repression. Purple lines represent functions of PIF1a-PIF1b heterodimers as deduced from this work.

Another conclusion based on the observed phenotypes is that PIF1 proteins may function as activators or repressors depending on the biological context. In Arabidopsis, the *pif1* mutant accumulates higher levels of carotenoids in the dark and produces more carotenoids and chlorophylls during de-etiolation (Huq et al., 2004; Moon et al., 2008; Toledo-Ortiz et al., 2010). In contrast, our results with tomato PIF1-defective mutants show only slightly increased levels of carotenoids in dark-grown seedlings but decreased levels of photosynthetic pigments in de-etiolating and light-grown seedlings (Fig. 2B). These results suggest that tomato PIF1a and PIF1b homologs might not be repressors but activators of photosynthetic development and leaf pigment biosynthesis (Fig. 5). Analysis of PIF1a-defective mutants of other Solanaceae species should provide valuable information on whether this is a general or species-specific phenotype.

Both PIF1a and PIF1b might also participate in the control of fruit production (i.e. total number of fruits per plant), as this parameter was reduced in single *pif1a* but not in double *pif1a pif1b* lines (Fig. 4A). We hypothesize that PIF1a-PIF1b heterodimers might activate fruit production whereas PIF1b-PIF1b homodimers might repress it (Fig. 5). The WT phenotype would result from a balance between activation and repression. Loss of PIF1a would lead to only repression (by PIF1b-PIF1b homodimers), whereas additional loss of PIF1b would remove both activation and repression pathways, resulting in a newly balanced situation and hence a WT phenotype in *pif1b* and *pif1a pif1b* plants. While other scenarios are possible, antagonistic roles of light signaling homologs are common. In Arabidopsis, PIF2 and PIF6 have antagonistic roles with PIF7 and PIFQ proteins for the control of light-triggered seedling de-etiolation (Pham et al., 2018). In the case of PIF2, it positively regulates seedling de-etiolation and photomorphogenesis by interacting with PIF1 and other PIFQ members, preventing them from regulating their target genes (Pham et al., 2018).

### PIF1a is the main PIF1 homolog regulating many other processes, including some that are not regulated by PIFs in Arabidopsis

Phenotypic analysis of the CRISPR/Cas9 mutants unveiled that there are many processes that would be regulated by just PIF1a. A striking case is the root hair elongation phenotype, which is not present in PIFQ-defective Arabidopsis mutants (Fig. 3). Arabidopsis phyB mutant lines were reported to develop longer root hairs (Reed et al., 1993), but it is possible that this phenotype depends on factors other than PIFs in Arabidopsis. Alternatively, scarcely explored PIFs (such as PIF2, PIF6 or PIF8) might have a major role in this process. Note that PIFs such as PIF4 and PIF5 have been identified as growth promoters in other organs, (Choi and Oh, 2016), notably hypocotyl elongation in response to shade (De Lucas et al., 2008), blue light (Pedmale et al., 2016) or temperature (Koini et al., 2009; Thines et al., 2014). Maybe during evolution, a neofunctionalization event took place and PIF1a acquired this elongation-promoting role specifically in root hairs (Fig. 5).

Flowering time also seems to be promoted by PIF1a, but not by PIF1b (Fig. 2D). Even though the differences between PIF1a-defective and WT lines are relatively small and are only observed when counting the number of leaves from germination to anthesis (but not when counting the number of days), they are statistically significant and consistent with the observation that PIF1 regulates flowering time in Arabidopsis (Wu et al., 2018) (Fig. 5). A number of other studies in the literature report differences in flowering when using one of the evaluated parameters but not when using the other (Calvert, 1959; Giliberto et al., 2005). Note that we grew our plants in the greenhouse under long day photoperiod, while domesticated tomato, in contrast with Arabidopsis, has been described as a day-neutral species (Soyk et al., 2017). Arabidopsis PIF4 and PIF5 were reported as thermosensory regulators of flowering (Kumar et al., 2012; Thines et al., 2014). Despite this, tomato PIF4 seems not to be related with this process, since the down-regulation of its expression leads to a reduction in flowers per truss, but not in flowering time (Rosado et al., 2019).

While seed germination, seedling de-etiolation, root hair development and flowering are biological processes present in both Arabidopsis and tomato, there are tomato-specific processes that are also regulated by PIF1a, such as those associated with fruit ripening. *PIF1a* is the only PIF-encoding gene that is up-regulated during fruit ripening (Rosado et al., 2016). One of the most characteristic features of tomato fruit ripening is the degradation of chlorophylls and the massive production of carotenoids when MG fruits start to ripe, changing their green color to red in the RR stage. PIF1a was first identified as a repressor of fruit carotenoid synthesis during ripening (Llorente et al., 2016b). In that work, an artificial microRNA approach was used to down-regulate *PIF1a* gene expression about 75%, resulting in ripe fruits that accumulated higher levels of total carotenoids (Llorente et al., 2016b). In contrast with those results, WT levels of carotenoids were detected in fruits from the *pif1a* mutant line generated in this work, which is predicted to be completely devoid of PIF1a (Supp Fig 7). An explanation for this (lack of) phenotype came from the careful analysis of off-target effects in the two studies that used gene knockdown approaches to conclude that PIF1a and PIF4 are repressors of carotenoid biosynthesis during fruit ripening (Llorente et al., 2016b; Rosado et al., 2019). In *PIF1a*-silenced fruits (Llorente et al., 2016b), the most strongly down-regulated off-targeted PIF was *PIF4* (~40%), while in *PIF4*-silenced fruits (Rosado et al., 2019) the most strongly down-regulated off-targeted PIF was *PIF1a* (~60%). Taking all these data together, we propose two possible explanations. First, PIF1a and PIF4 might be functionally redundant and play a similar role in modulating fruit carotenoid biosynthesis during ripening. This hypothesis involves that higher carotenoid contents would only be observed when both PIF1a and PIF4 are down-regulated. Because *PIF4* expression is not affected in the *pif1a* mutant generated in this work (Supp Fig 5), PIF4 levels would be high enough to repress carotenoid overaccumulation in the *pif1a* line. The second hypothesis states that PIF4, but not PIF1a, would be involved in repressing carotenoid production during ripening. Thus, reduced *PIF4* transcript levels in fruits of knock-down *PIF1a* and *PIF4* lines, but not in our CRISPR-Cas9 mutants, would lead to higher carotenoid contents.

Fruit growth in terms of size (Fig. 4B) and weight (Fig. 4C) increased in *pif1a* but also in the *pif1a pif1b* double mutant. Based on the results of the fruit desiccation experiment (Supp Fig. 8) we concluded that the difference in size might be due to a difference in tissue mass, not in water content of the fruits, pointing out PIF1a as a repressor of fruit growth (Fig. 5). Arabidopsis PIF1 has been shown to regulate the expression of cell-wall-related genes (Oh et al., 2009; Shi et al., 2013). Alteration of cell-wall structure by PIF1a might explain the fruit softening phenotype of PIF1a-defective fruits (Fig 4D). Thus, PIF1a might be involved in the loosening of the cell-wall to allow root hair elongation and fruit softening while repressing fruit growth by a different mechanism (Fig. 5). This contrasting and opposite role of PIF1a in different tissues and developmental stages can be due to different interactions with different partners depending on the tissue and the developmental stage. For instance, protein-protein interaction between PIFs and with other factors can modulate their capacity to bind to the DNA (Pham et al., 2018). In this way, PIF1a might have specific interactors to promote root hair elongation or ripe fruit softening, while a different set of interactors would repress fruit growth.

Arabidopsis PIF4 has an important role in promoting hypocotyl elongation in response to different light and temperature cues (Koini et al., 2009; Choi and Oh, 2016; Pham et al., 2018). Supporting the conclusion that PIF4 has conserved a growth-promoting role in different plant species and tissues, tomato knock-down lines for PIF4 develop smaller fruits (Rosado et al., 2019). Therefore, PIF1a and PIF4 appear to play antagonistic roles in the regulation of fruit size. A closer look at the expression patterns of the genes encoding PIF1a and PIF4 during fruit development shows that the expression of *PIF1a* is low at the early stages of fruit development (when growth takes place by cell division and proliferation) and slowly increases as cells expand (until the MG stage) and fruits begin to ripe, peaking at the RR stage (Supp Fig 9). In contrast, PIF4 expression peaks at the MG stage (i.e. when fruit reaches its final size) and then drops during ripening (Supp Fig 8). Based on these data, we speculate that PIF4 is the main promoter of fruit growth. Similar to that discussed above on the opposite role of PIF1a-PIF1b heterodimers and PIF1b homodimers for the regulation of fruit production (Fig. 5), it is possible that PIF1a-PIF4 heterodimers could antagonize the activating role of PIF4, perhaps by preventing binding of PIF4 homodimers to fruit-promoting target genes. As PIF1a levels increase during fruit development, heterodimers become more abundant and PIF4 homodimers decrease, resulting in a progressive attenuation of growth. Then, after the MG stage, PIF4 levels drop and growth stops. The peak of PIF1a expression at the RR stage is most likely unrelated to growth but required for fruit softening. Whether tomato PIF4 also interacts with PIF1a to regulate root hair elongation remains unknown.

### Evolutionary implications of PIF1 duplication

As a summary of this work, graphically represented in Fig. 5, we conclude that some of the PIF1-regulated processes in Arabidopsis are also regulated by PIF1 homologs in tomato (including seed germination, photosynthetic pigment accumulation and flowering time). A second group includes processes that are regulated by PIF1 homologs in tomato but not by PIF1 in Arabidopsis. From them, some occur in both plant species (such as root hair elongation) but others are specific of tomato (fruit ripening). Moreover, the duplication of PIF1 present in tomato led to some of the above-mentioned processes being regulated by both PIF1a and PIF1b (seed germination, photosynthetic pigment accumulation and fruit production) while the rest became controlled just by PIF1a. Interestingly, we did not find any process that would be regulated exclusively by PIF1b (Fig. 5).

The duplication of *PIF1* has been analyzed in detail previously (Rosado et al., 2016). The duplication of *PIF1* in *PIF1a* and *PIF1b* is predicted to have happened just before the divergence between tomato and potato (Rosado et al., 2016; Sato et al., 2012). We hypothesize that before the divergence between Arabidopsis and *Solanum* there were likely some processes that were already regulated by PIF1 homologs in both species (e.g., seed germination, photosynthetic pigment contents or flowering time). During the following millions of years, the duplication of PIF1 in tomato led to newly acquired functions. While PIF1a remained as the main isoform, the presence of PIF1b provided robustness to essential processes (e.g. seed germination) and allowed to regulate new processes via heterodimerization (e.g. fruit production). On the other hand, some pre-existing processes regulated by PIF1 in Arabidopsis became controlled just by PIF1a in tomato, such as flowering time. The reason behind this might have been the mutation identified in PIF1b that leads to the loss of interaction with phyB. PIF1a also acquired the capacity to regulate new processes. The root hair elongation phenotype is a striking example with two possible explanations. The first one is a neofunctionalization process, by which PIF1a ended up involved in the regulation of the elongation of this cell type. A second possibility is that the original PIF1 was controlling this trait, but during evolution this role was lost in Arabidopsis but remained in tomato. In fruit, PIF1a and, likely, PIF4 might have re-adapted their already existing role in the regulation of cell elongation (Paik et al., 2017) to determine the final fruit size. There are several examples of transcription factors that have been recycled during evolution to regulate new processes, like fruit ripening (Gapper et al., 2013). For instance, in Arabidopsis SHATTERPROOF 1 (SHP1) and SHP2 are important regulators of valve margin identity and its subsequent dehiscence zone (Liljegren et al., 2000). By contrast, their closest homolog in tomato, TOMATO AGAMOUSLIKE1 (TAGL1) controls fleshy fruit expansion (Itkin et al., 2009; Vrebalov et al., 2009; Gimenez et al., 2010). Another example is SEPALLATA 4 (SEP4), which in Arabidopsis regulates organ identity in flowers (Ditta et al., 2004). The closest homolog to SEP4 in tomato in RIPENING INHIBITOR (RIN), which is a key regulator of fruit ripening and controls climacteric respiration and ethylene biosynthesis (Fujisawa et al., 2011; Martel et al., 2011). Our results suggest that processes of specialization and neofunctionalization besides those detected here by analyzing the phenotypes of CRISPR-Cas9 mutants would have taken place during evolution after the splitting of Arabidopsis and *Solanum* groups. Further approaches, such as transcriptomics or metabolomic analysis, would provide some clues about which new tomato processes became regulated by PIF1a and PIF1b.

## ACKNOWLEDGEMENTS

We thank greatly Jordi Gine-Bordanaba for assistance with the texture analyzer and M. Rosa Rodriguez-Goberna for technical help. The work was funded by Spanish grants BIO2017-84041-P and PID2020-115810GB-I00 from the Agencia Estatal de Investigacion (AEI) and FEDER and 202040E299 from Consejo Superior de Investigaciones Científicas (CSIC) to MRC. We also acknowledge the AEI support for the PRIMA project UToPIQ (PCI2021-121941) and the Generalitat Valenciana grant PROMETEU/2021/056. MSM was supported by PhD fellowship BES-2015-072725 from the Spanish Ministerio de Economia y Competitividad. M.V.B. was funded with a Spanish Ministry of Education, Culture and Sports PhD fellowship (FPU14/05142). L.M. received a predoctoral fellowships from La Caixa Foundation (INPhINIT fellowship LCF/BQ/IN18/11660004) which received funding from the EU-H2020 through MSCA Grant 713673.

## MATERIALS AND METHODS

### Plant material and growth conditions

*Nicotiana benthamiana* and tomato plants were grown under standard long day greenhouse conditions (14 h light at 27 ± 1°C and 10 h dark at 24 ± 1°C). Tomato plants were transformed as previously described (Fernandez et al., 2009). Seed germination assays were performed as was previously described (Appenroth et al., 2006). Briefly, Tomato seeds were surface-sterilized for 10 min with 40% (v/v) NaClO, washed 3 times with sterile water and sown on sterile Murashige and Skoog (MS) medium containing 1% agar and no sucrose. After that, seeds were incubated in dark conditions and exposed hourly to 5-min R (200 μmol m^-2^ s^-1^) or FR (100 μmol m^-2^ s^-1^). Temperature was always kept at 25 °C. Radicle protrusion was used as a criterion for judging seed germination. For de-etiolation experiments, tomato seeds were surface-sterilized for 10 min with 40% (v/v) NaClO, washed 3 times with sterile water and sown on sterile wet cotton. After that, seed were incubated in dark conditions, except the Light control of the experiment, that was incubated in continuous light conditions. After 7 days, the seedlings were transferred to light. Samples were collected at the indicated time points after exposure to light. Temperature was always kept at 25 °C. Flowering time was assessed as was previously reported in tomato (Dielen et al., 1998). Basically, after sowing the seeds we waited until the first flower reached the anthesis stage. In that moment we measure two parameters: The number of leaves and the days post-sowing when the anthesis took place. Fruit production was measured by counting the number of produced fruits per plants in 19-weeks-old plants in greenhouse conditions. The volume of the fruits was assessed by measuring the displaced volume in a graduated cylinder of a group of 10 representative fruits. The weight of the fruits was assessed by two different methods. First, 100 fruits were weight individually. Second, 15 fruits were weight together as a group. This second method was used to compare the weight difference between fresh samples and dry samples. To dry the fruits, all the 15-fruits groups were incubated for 4 days at 90 °C. Fruit hardness was measured with a texture analyzer (Stable Micro Systems, TA-XT2i) fitted with 50 mm plate probe to perform a compression test (Kabas and Ozmerzi, 2008; Camps and Gilli, 2017). The instrument was set to measure the mechanical work needed to reach 5% deformation of the original form of the fruit.

### Transient expression assays

Leaves from 4-week-old *N. benthamiana* plants were entirely infiltrated with the desired combination of *Agrobatoerium tumefaciens* strains in the infiltration buffer (10 mM MES pH5.5.-6, 10 mM MgSO4, acetosyringone 100 μM). After agroinfiltration, the plants were left in the greenhouse for 3 days. Leaves were used to perform confocal microscopy analysis. In the case of BiFC experiments, equal volumes of Agrobacterium cultures were mixed to get the indicated combinations in Fig. 1D and Supp Fig 3.

### DNA constructs and genotyping

The coding sequences of PIF1a and PIF1b were cloned into a gateway ENTRY vector (pDNOR207, Thermo Fisher). After that, using a gateway LR reaction (Thermo Fisher), the CDSs were transferred to the final expression vectors, in frame with the GFP gene for subcellular localization (pDONR221 and pK7FWG2), and with N- or C-terminal sequence of GFP for BiFC experiments (pYFN43 and pYFC43 respectively) (Belda-Palazón et al., 2012). For the CRISPR/Cas9 constructs two guide RNAs (sgRNA) were cloned via cut-ligation reaction with BbsI (Thermo) and T4 DNA ligase (Thermo) in a Gateway ENTRY sgRNA shuttle vector (Ritter et al., 2017). Next, using a gateway LR reaction (Thermo Fisher), the two sgRNA modules were then combined with pDE-Cas9Km vector (Ritter et al., 2017) to yield the final expression clone. First identification of CRISPR/Cas9 mutations was assessed with the TIDE (Tracking of Indels by DEcomposition) web tool. We analyze the chromatograms individually to identify the short insertion or deletion, using as template a chromatogram of a WT plant. The alignment was performed in Benchling (www.benchling.com) using MAFFT as an alignment program. Once the insertions or deletions were identified, CRISPR/Cas9 lines were genotyped using a PCR to amplify 500 bp of the mutated version. During the identification of the mutation we detected that the mutation in PIF1a removed a restriction site for DrdI and the mutation in PIF1b removed a restriction site for NlaIV. The amplified fragments were then digested with the indicated enzymes.

### Gene expression profiling

Expression data of different tissues were retrieved from the Bio-Analytic Resource for Plant Biology (BAR, University of Toronto), using the tomato tool available in that website and PIF1a to PIF1b Solyc IDs. Co-expression analysis was performed as previously described (Ahrazem et al., 2018). Briefly, tomato PIF1a to PIF1b Solyc IDs were used to retrieve all the expression data available for different cultivars/tissues/treatment in the Tomexpress database. Subsequently, global gene coexpression (GCN) analysis was carried out by calculating pairwise Pearson correlation coefficients between each PIF gene against the tomato genome was computed, and Fisher’s Z–transformation was used to test the statistical significance of the pairwise correlations.

### Quantification of metabolites

Carotenoids, chlorophylls, tocopherols were extracted as described (Barja et al., 2021). Ethylene production in tomato fruit was assessed by adapting a previous method (Pereira et al., 2017). Specifically, we collected tomato fruits at MG stage and incubated them in long day greenhouse conditions in opened 50-mL tubes. Once a day, the tubes were sealed with Parafilm (BEMIS) and incubated for one hour. After that hour, 30 mL of the headspace air was collected by injecting a syringe through the Parafilm layer. That samples were finally collected in a GC-MS tube and analyzed. GC-MS analysis was performed with an Agilent 7890A gas chromatograph coupled to a 5975C mass selective detector.

### Confocal and electron microscopy

Subcellular localization, protein stability and BiFC experiments were determined by analyzing agroinfiltrated leaf samples with an Olympus FV 1000 confocal laser-scanning microscope. GFP signal and chlorophyll autofluorescence were detected using an argon laser for excitation (at 488 nm) and a 500–510 nm filter for detection of GFP fluorescence and a 610–700 nm filter for detection of chlorophyll fluorescence. In the case of protein stability, same conditions were used strictly to take all the images. First, a field of 20 – 30 nuclei was photographed. After that, the sample was incubated during 30 min in Red (200 μmol m-2 s-1) and Far Red light (100 μmol m-2 s-1). The exact same field was photographed again after that exposure, using the same parameters as before. The intensity of the fluorescence signal was measures using the ImageJ software. Individual Regions Of Interest (ROI) were created for each nucleus and identical-size ROI were used to compare the same nucleus before and after the light exposure. The intensity of fluorescence was measured in each pixel of all the ROI and each ROI was considered an individual replicate within one population.

Root samples were washed 2 washes on 10 min in absolute ethanol and after that they were dry by critical point using an equipment Emitech K850. Then, the samples were mounted on the supports of the SEM suing a biadhesive conductive disk and coated with a thin layer of gold (about 60nm) to make them conductive electrical. This was done with the SEM coating system model SC510 (Fisons Instruments). The study was carried out using a Field Emission Scanning Electron Microscope J-7001F (Jeol) using a secondary electron detector to analyze the topology of the samples.

