## Supplemental Data for "Conserved and newly acquired roles of PIF1 homologs in tomato (*Solanum lycopersicum*)"

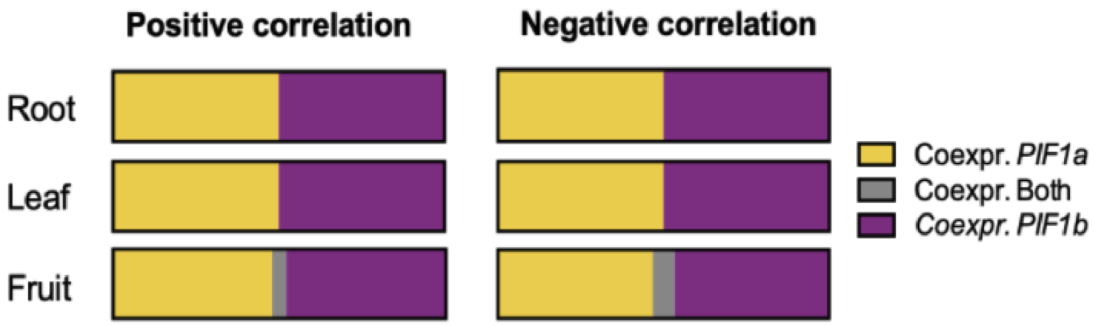


**Supplemental Figure 1.** Distribution of the 500 most co-expressed genes with PIF1a (yellow) and PIF1b (purple) in roots, leaves and fruits. Overlapping genes co-expressed with both PIF1a and PIF1b are represented in grey.

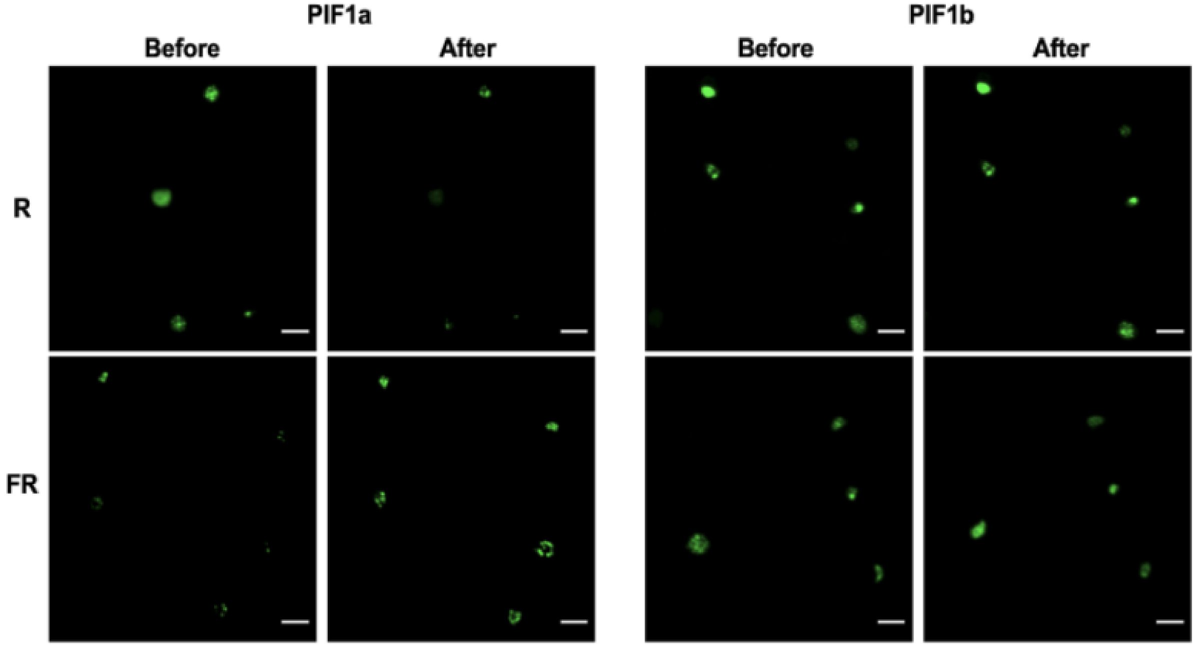


**Supplemental Figure 2.** Confocal microscopy images of leaf areas expressing PIF1a or PIF1b fused to GFP, before and after exposure to either R of FR for 30 min. Pictures show the same area of the leaf with the same laser conditions. Scale bar = 20 μm.

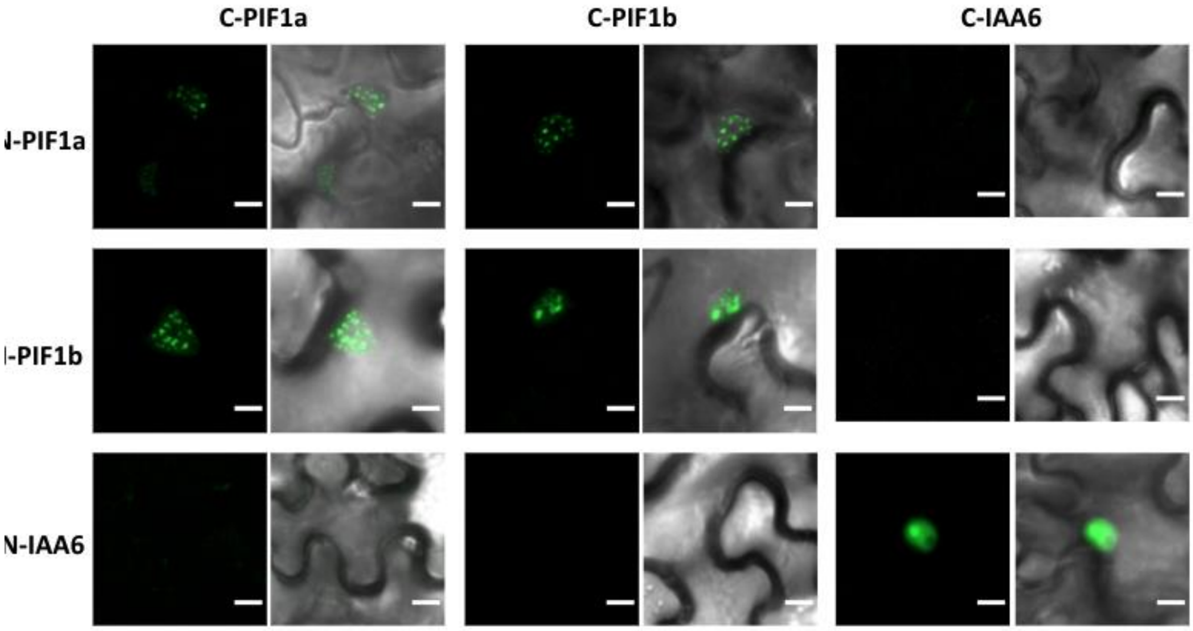


**Supplemental Figure 3.** BiFC analysis of PIF1a and PIF1b protein-protein interaction.Confocal microscopy images of GFP fluorescence in N. benthamiana leaf cells transiently expressing the indicated proteins fused to N- or C-terminal GFP halves for BiFC analysis. Images of representative nuclei showing either GFP fluorescence alone (left) or overlapped with bright field images (right) are shown for every combination. Scale bar = 10 μm.

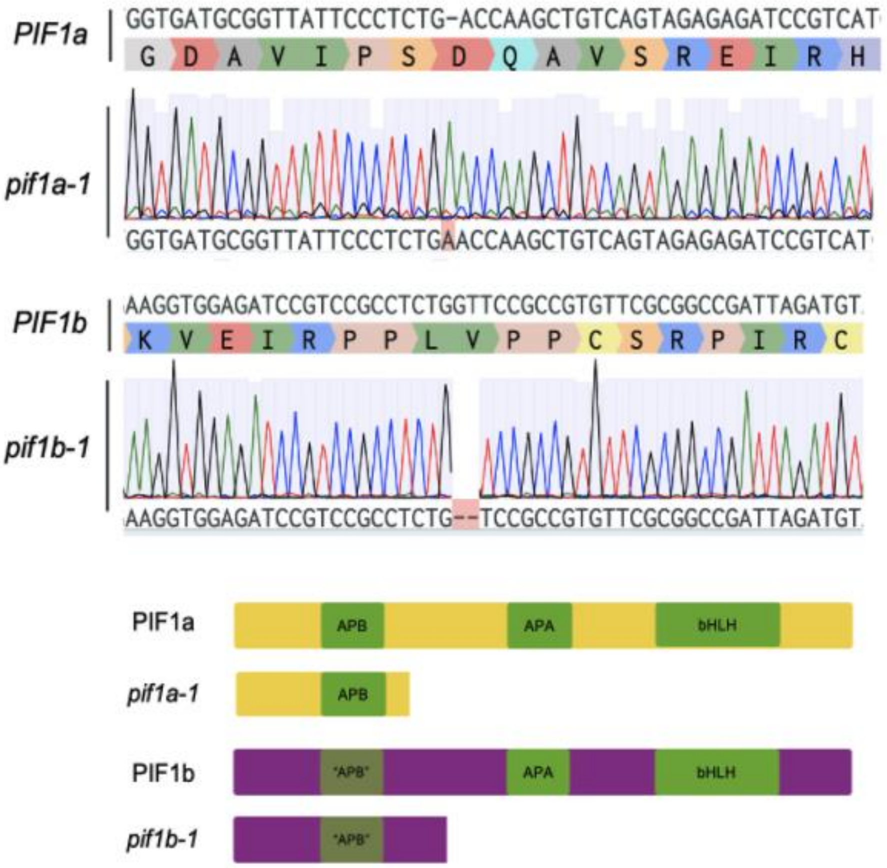


**Supplemental Figure 4.** Analysis of CRISPR/Cas9 mutations in PIF1a and PIF1b. Upper: Chromatograms of the sequences analyzed by the TIDE platform. Indels in the identified alleles are highlighted. Lower: Schematic representation of PIF1a and PIF1b proteins in the CRISPR/Cas9-generated mutant alleles.

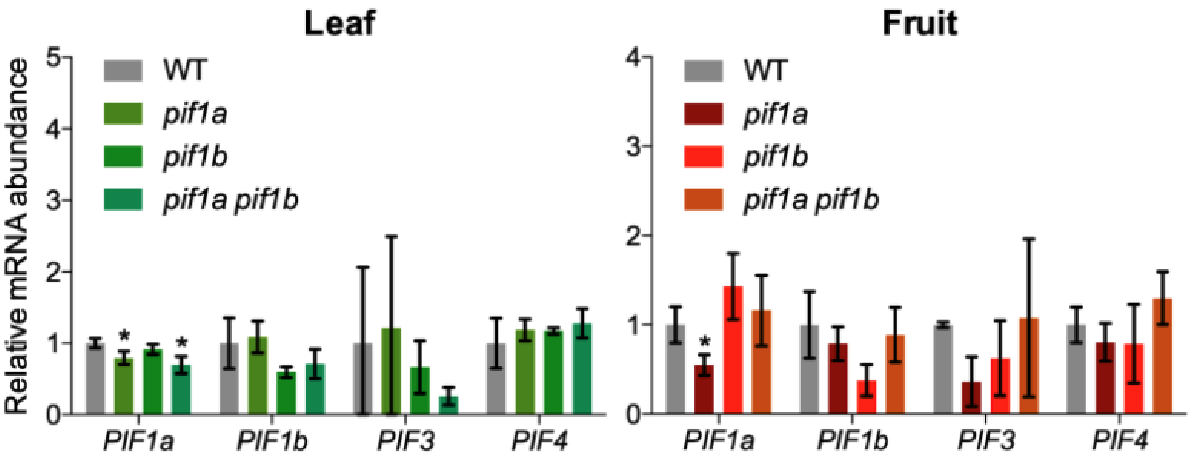


**Supplemental Figure 5.** qPCR analysis of PIF-encoding transcripts in mature leaves and MG fruit from tomato CRISPR-CAs9 mutants defective in PIF1a, PIF1b or both. Error bars indicate SD of 3 biological replicates in the indicated tissues. Asterisks mark statistically significant changes in student's t test (* = p < 0.05).

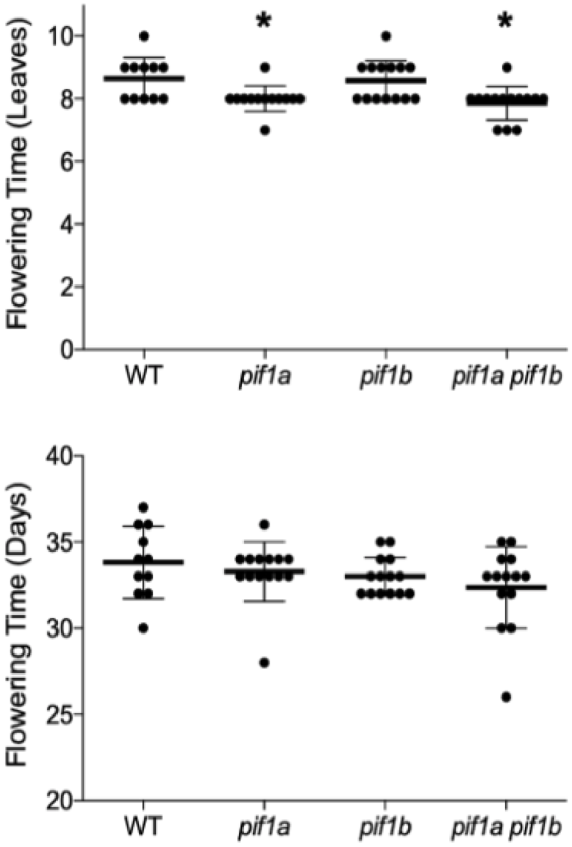


**Supplemental Figure 6.** Flowering time in tomato pif1 mutants. Plants were grown on soil in the greenhouse under long day conditions. Error bars indicate SD of at least 11 different plants. Asterisks mark statistically significant changes in the indicated genotype compared to the WT according to one-way ANOVA (* = $p < 0.05$).

Upper. Number of leaves in the plant when the first flower reached anthesis.

Lower. Number of days from sowing until the first flower reached anthesis.

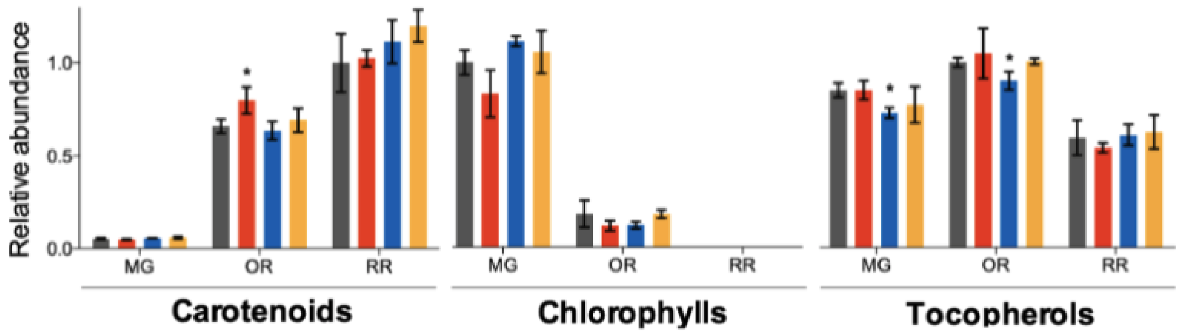


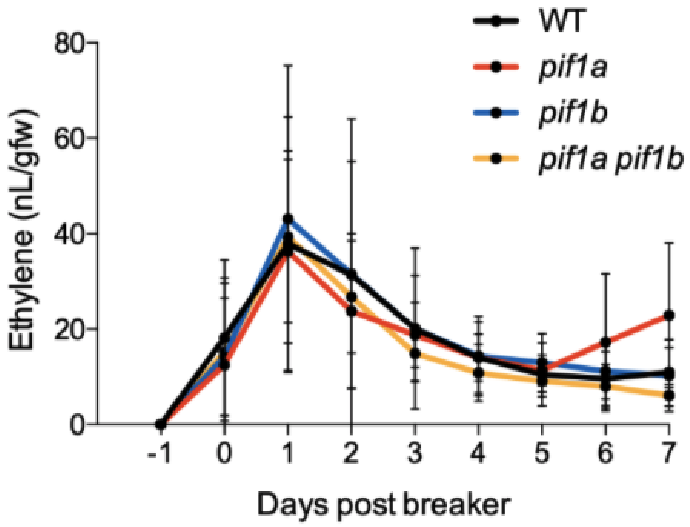


**Supplemental Figure 7.**

Upper. Isoprenoid levels in fruits of tomato pif1 mutants at different stages of ripening. The indicated metabolites were quantified in 3 different pools of 4 fruits from plants grown on soil in the greenhouse under long day conditions. Error bars indicate SD of the 3 biological replicates. Asterisks mark statistically significant changes in the indicated genotype compared to the WT according to t-student test (* = $p < 0.05$).

Lower. Ethylene levels during fruit ripening in tomato pif1 mutants. Fruits from plants grown on soil in the greenhouse under long day conditions were harvested at the MG stage and transferred to sealed 50 mL tubes (one fruit per tube). Ethylene was quantified every day from the headspace inside the tube after one hour of incubation. Time was set at 0 when fruits reached the breaker (BR) stage. The graph plots results from one day before BR to 7 days after BR. Error bars indicate SD of at least 5 different fruits per timepoint.

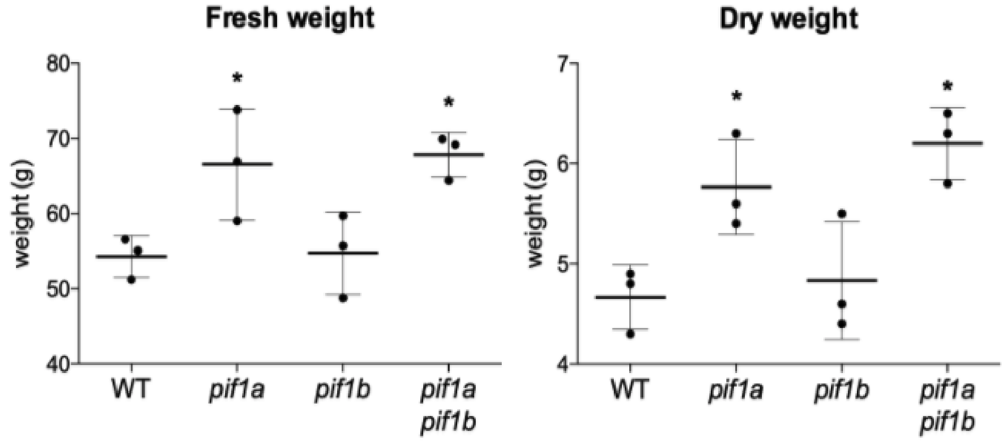


**Supplemental Figure 8.** Weight of 3 groups of 15 RR fruits before (fresh) and after (dry) incubation in an oven until complete loss of water.Error bars indicate SD. Asterisks mark statistically significant changes in the indicated genotype compared with WT according to one-way ANOVA (* = $p < 0.05$)

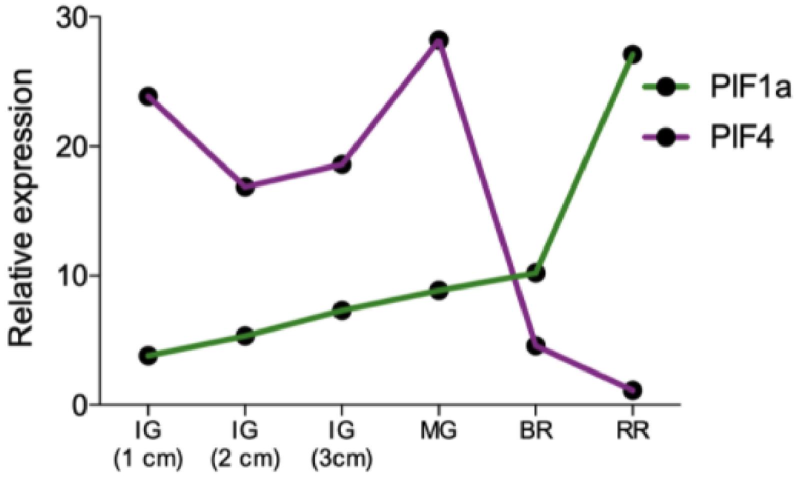


**Supplemental Figure 9.** Expression profiles of tomato PIF1a and PIF4 during fruit development and ripening. Data obtained from The Bio-Analytic Resource from Plant Biology (BAR, University of Toronto).
